# Cell-Type Specific Profiling of Histone Post-Translational Modifications in the Adult Mouse Striatum

**DOI:** 10.1101/2022.01.17.476614

**Authors:** Marco D. Carpenter, Delaney K. Fischer, Shuo Zhang, Allison M Bond, Kyle S. Czarnecki, Morgan T. Woolf, Hongjun Song, Elizabeth A. Heller

## Abstract

Histone post-translational modifications (hPTMs) regulate gene expression via changes in chromatin accessibility and transcription factor recruitment. At a given locus, the coordinated enrichment of several distinct hPTMs regulate gene expression in response to external stimuli. However, neuronal hPTMs have been primarily characterized in bulk brain tissue and/or tissue pooled across subjects. This obscures both cell-type and individual variability, features essential to understand individual susceptibility to psychiatric disease. To address this limitation, we optimized a hybrid protocol, ICuRuS, to profile both activating and repressive hPTMs in neuronal subtypes from a single mouse. We report here profiling of striatal medium spiny neuron (MSN) subtypes, genetically defined by expression of Adenosine 2a Receptor (A2a) or Dopamine Receptor D1 (D1), which differentially regulate reward processing and pathophysiology. Using ICuRuS, we defined genome-wide, A2a- or D1-specific combinatorial hPTM profiles, and discovered regulatory epigenomic features at genes implicated in neurobiological function and disease.

## Main

Epigenetic regulation in the heterogenous brain remains challenging to decipher. Bulk tissue analysis reflects the average of cell-type specific changes, which impedes the interpretability of causal relationships between epigenetic modifications, regulation of gene expression and behavior ^1–3^. Cell-type specific RNA-seq allows for the deconvolution of bulk tissue RNA-seq data, but cell-type specific epigenomic profiling in brain is only emerging ^4–7^. Fortunately, recent methodological advances facilitate cell-type specific epigenetic profiling in brain ^8–11^. Given that pathogenic gene expression results from cell-type specific abnormalities ^12–15^, a specific cell-type may contain intrinsic disease-causing factors or be one of many cell-types affected by or causing a common pathology. For example, in Parkinson’s disease, loss of dopaminergic neurons leads to perturbations in function of several additional cell-types, including MSNs ^12^. In contrast, genes implicated in schizophrenia are expressed specifically in MSNs rather than interneurons, astrocytes, or glia ^13,16^. Promisingly, many schizophrenia-related genes are targets of anti-psychotic medications ^13,14^. With respect to substance use disorder (SUD), a robust literature on A2a and D1 specific connectivity ^17,18^, gene expression ^19^ and chromatin accessibility ^20^, supports the notion that MSN-subtype specific function and epigenetic regulation underlies reward pathophysiology. However, MSN-specific hPTM profiling from a single animal has not yet been reported.

Chromatin immunoprecipitation sequencing (ChIP-seq) has been widely used to profile hPTMs in specific neuronal populations but this method requires large numbers of nuclei and is prone to high background ^12–15,21,22^. We developed ICuRuS by combining methods for isolation of nuclei tagged in specific cell-types ^9^ (INTACT) and hPTM profiling by <u>c</u>leavage <u>u</u>nder targets & <u>r</u>elease <u>u</u>sing nuclease ^23,24^ (CnR), followed by next generation <u>s</u>equencing. This approach was developed to address the following limitations of ChIP-seq: (1) At low starting material, such as one specific neuronal subtype, the reliability and detection sensitivity of subtle, physiologically-relevant changes is greatly diminished ^21,22^. This necessitates pooling across subjects, which hinders downstream correlations between hPTMs and individual subject behavior and/or pathology ^5^. (2) The integrity of epigenomic features are impacted by the nuclear isolation technique ^8,9,25,26^. For example, fluorescence activated cell sorting (FACS) leads to cellular stress, artifactual DNA shearing, and increased sequencing noise ^23^. To compensate for high noise, data is normalized to input chromatin, requiring greater sequencing costs. (3) ChIP often requires formaldehyde fixation, which can reduce the efficiency of immunoprecipitation by masking protein epitopes and stabilize highly transient chromatin binding, leading to false positive reads ^27–29^. ChIP can be conducted without cross-linking, but problems still remain, such as insufficient binding efficiency ^23^. ICuRuS overcomes these obstacles and allows for low-cost sequencing with limited starting material.

Here, we characterized cell-type specific hPTM profiles in striatal MSNs. First, we isolated A2a and D1 nuclei of sufficient number, purity, and quality for epigenomic profiling. To model and predict cell-type specific gene expression patterns, we profiled both activating and repressive hPTMs, H3K4me3 (histone H3 lysine 4 tri-methylation) and H3K27me3, respectively. These hPTMs are important regulators of poised and activity-dependent transcription, mechanisms implicated in these cell types ^30–33^. We showed A2a and D1 nuclei differ in their relative enrichment of H3K4me3 and H3K27me3 at cell-type specific genes but are largely similar in overall pattern of hPTM enrichment across the genome. Specifically, analysis of genes enriched in both H3K4me3 and H3K27me3 revealed that cell-type specific gene expression is associated with either enrichment or depletion of H3K4me3 or H3K27me3, with or without changes in the opposing modification. We highlighted our findings on *Nr4a1*, a gene that we and others have characterized as particularly relevant to cocaine use disorder and rapidly activated in response to cocaine exposure ^33–36^. We report here that *Nr4a1* is enriched in H3K4me3 and depleted in H3K27me3 in both MSN subtypes ^36^. Overall, these data highlight that high resolution epigenomic profiles generated by ICuRuS defined combinatorial relationships between neuronal-subtype specific hPTMs and gene expression in the mammalian brain.

## Results

### INTACT purified D1 and A2a MSN nuclei from a single mouse brain

We employed INTACT to isolate A2a and D1 nuclei at sufficient number and quality for epigenomic profiling. To generate a mouse line for affinity purification of striatal A2a and D1 nuclei, we crossed the established SUN1-sfGFP-Myc mouse line ^9^, expressing a GFP-affinity-tagged SUN1 nuclear receptor under the control of a loxP-3xPolyA-loxP transcriptional stop cassette, to either A2a- or D1-Cre mouse lines. A2a-Cre; or D1-Cre; SUN1-GFP mice are healthy, fertile and display no phenotypic abnormalities (**Supplementary Figure 1A**). Double immunohistochemistry of GFP and A2a or GFP and D1 showed expression of nuclear SUN1-GFP in the target cell-type (**Figure 1A, B**). Using INTACT ^9^, we isolated A2a or D1 nuclei from striatum of a single mouse with an anti-GFP antibody (**Figure 1C, D**). Fluorescence microscopy showed a sufficient number of nuclei (8-10,000) were recovered for downstream hPTM profiling by CnR (**Figure 1E**) ^23,24,37–39^ and gene expression profiling using qPCR of high quality mRNA (**Supplementary Figure 1B**). To determine the specificity of the isolated nuclei, we measured expression cell-type specific genes, *A2a* and *Drd1*, in the affinity purified and ‘flow-through’ fractions after INTACT of each cell-type **(Figure 1F-G)**. Following A2a INTACT, A2a mRNA was enriched and Drd1 mRNA was depleted in the A2a affinity purified fraction relative to the flow-through (**Figure 1F**). Following D1 INTACT, Drd1 mRNA was enriched and A2a mRNA was depleted in the D1 affinity purified fraction relative to the flow-through (**Figure 1G**). Note that A2a mRNA was undetectable in D1 nuclei and Drd1 mRNA was >20-fold depleted in A2a isolated nuclei (data not shown). These data show that INTACT successfully isolated A2a and D1 nuclei from a single mouse striatum.

**Figure 1.**
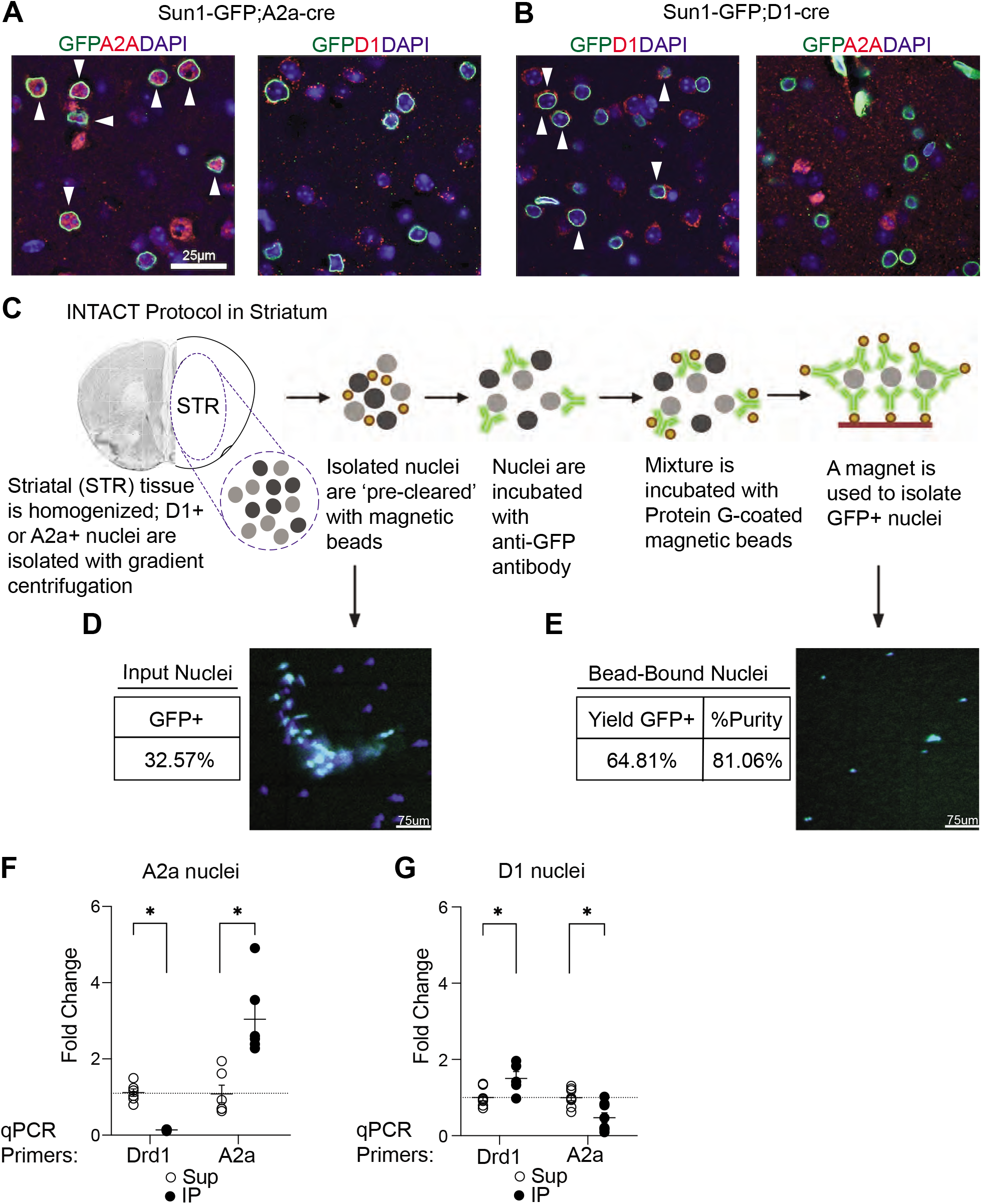
SUN1-GFP facilitates isolation of D1 or A2a neurons in the Striatum. **A)** Immunohistochemistry displaying DAPI/GFP/A2A (left) and D1 (right) from the striatum of SUN1-GFP;A2a-CRE+ animals. Arrows indicate co-localization of all three markers. **B)** Immunohistochemistry displaying DAPI/GFP/DRD1 (left) and A2A (right) from the striatum of SUN1-GFP; D1-CRE+ animals. Arrows indicate co-localization of all three markers. **C)** INTACT schematic created at Biorender.com. **D)** Purity (&) of Cre+ cells in STR. Image depicts total nuclei isolation. **E)** Yield (&) and Purity (&) of GFP+ nuclei. Image depicts GFP+ nuclei isolation. **F)** mRNA validation for INTACT Sun1-GFP; A2a-Cre+ vs bulk (Cre-nuclei) **G)**mRNA validation for INTACT Sun1-GFP;D1-Cre + vs bulk (Cre-nuclei) (right).

### Epigenomic profiling using CnR in N2a Cells

We first validated H3K4me3 and H3K27me3 CnR in N2a cells ^40^ **(Supplemental Figure 2A-J)** finding that H3K4me3 and H3K27me3 profiles aligned with published N2a ChIP-seq data ^41,42^ (**Supplemental Figure 2B)**. For H3K4me3 and H3K27me3, the genomic distribution of reads across replicates was highly similar, based on Pearson’s correlation coefficient (PCC; H3K4me3 PCC: 0.99, H3K27me3 PCC: 0.94; **Supplemental Figure 2C, D**), and correlated with corresponding ChIP-seq data for each modification (H3K4me3 PCC: 0.78, H3K27me3 PCC: 0.63; **Supplemental Figure 2C, D**), but not with IgG control (PCC < 0.1; **Supplemental Figure 2C, D**). Moreover, H3K4me3 and H3K27me3 CnR reads were enriched around peaks called from corresponding N2a ChIP-seq data ^41,42^ (**Supplemental Figure 2E, F**), indicating that CnR recapitulated features of ChIP-seq. Quantification of the fraction of reads in peaks (FRiP) revealed that H3K4me3 CnR in N2a cells produces high quality data with negligible signal to noise, similar to published ChIP-seq FRiP in this cell line **(Supplemental Figure 2G)**. H3K27me3 CnR FRiP also showed high signal to noise which was slightly enhanced compared to published ChIP-seq FRiP **(Supplemental Figure 2I)**. Additionally, there was a 63& and 35& overlap in H3K4me3 and H3K27me3 peaks, respectively, between N2a CnR and previously published ChIP-seq data ^42^ (**Supplemental Figure H, J)**. Thus, CnR successfully profiled H3K4me3 and H3K27me3 modifications in N2a cells.

### Epigenomic profiling of MSNs using ICuRuS is robust and reproducible

We next optimized ICuRuS by combining <u>I</u>NTACT and <u>CU</u>T&<u>RU</u>N-<u>s</u>eq, to profile hPTMs in specific MSN subtypes of the mouse striatum. CnR is easily applied to neuronal nuclei following INTACT, as sfGFP-SUN1+ nuclei are immobilized during INTACT on paramagnetic beads ^23,24^. Bead-bound nuclei were then incubated with antibodies against H3K4me3 and H3K27me3 and subjected to antibody guided nucleosomal MNase cleavage **(see Supplemental Figure 2A)**, followed by NGS and validation by comparison to published ChIP-seq datasets. We found that A2a and D1 CnR H3K4me3 profiles were similar to NAc H3K4me3 ChIP-seq profiles ^43^ **(Figure 2A)**. To further validate striatal ICuRuS, we computed correlation matrices comparing A2a and D1 ICuRuS replicates to NAc ChIP-seq data using read coverages for the entire genome in 1-kb bins ^43^. Within A2a and D1 cell-types, H3K4me3 ICuRuS replicates were highly similar (PCC: 0.99 for both A2a and D1; **Figure 2B**). Between A2a and D1 cell-types, H3K4me3 replicates were more similar to each other (PCC: 0.97) than to bulk striatal ChIP-Seq (PCC: 0.53-0.73; **Figure 2B**). Within A2a and D1 cell-types, H3K27me3 ICuRuS replicates were highly similar (A2a PCC: 0.89; D1 PCC: .76; **Figure 2C**). Between A2a and D1 cell-types, H3K27me3 ICuRuS replicates were more similar to each other (PCC: 0.76 – 0.77) than to bulk striatal ChIP-Seq (PCC: 0.37-0.48; **Figure 2B**). Next, we quantified H3K4me3 and H3K27me3 ICuRuS signal from each cell-type in peaks called from bulk NAc ChIP-seq ^43^. We found that both A2a and D1 ICuRuS H3K4me3 and H3K27me3 signal was enriched at ChIP-seq peak centers (**Figure 2D, E**). In contrast, control ICuRuS using IgG or no antibody resulted in sparse enrichment around the peaks (**Figure 2D, E; Supplemental Figure 2K-P**). We next called ICuRuS peaks and found 78& and 42& of H3K4me3 and H3K27me3 peaks overlapped, respectively, between ICuRuS and previously published ChIP-seq data ^43^ **(Supplemental Figure 3A, B; Supplementary File 1)** and A2a and D1 ICuRuS H3K4me3 and H3K27me3 reads populated corresponding NAc ChIP-Seq peaks at comparable levels (**Supplemental Figure 3C, D)**.

**Figure 2.**
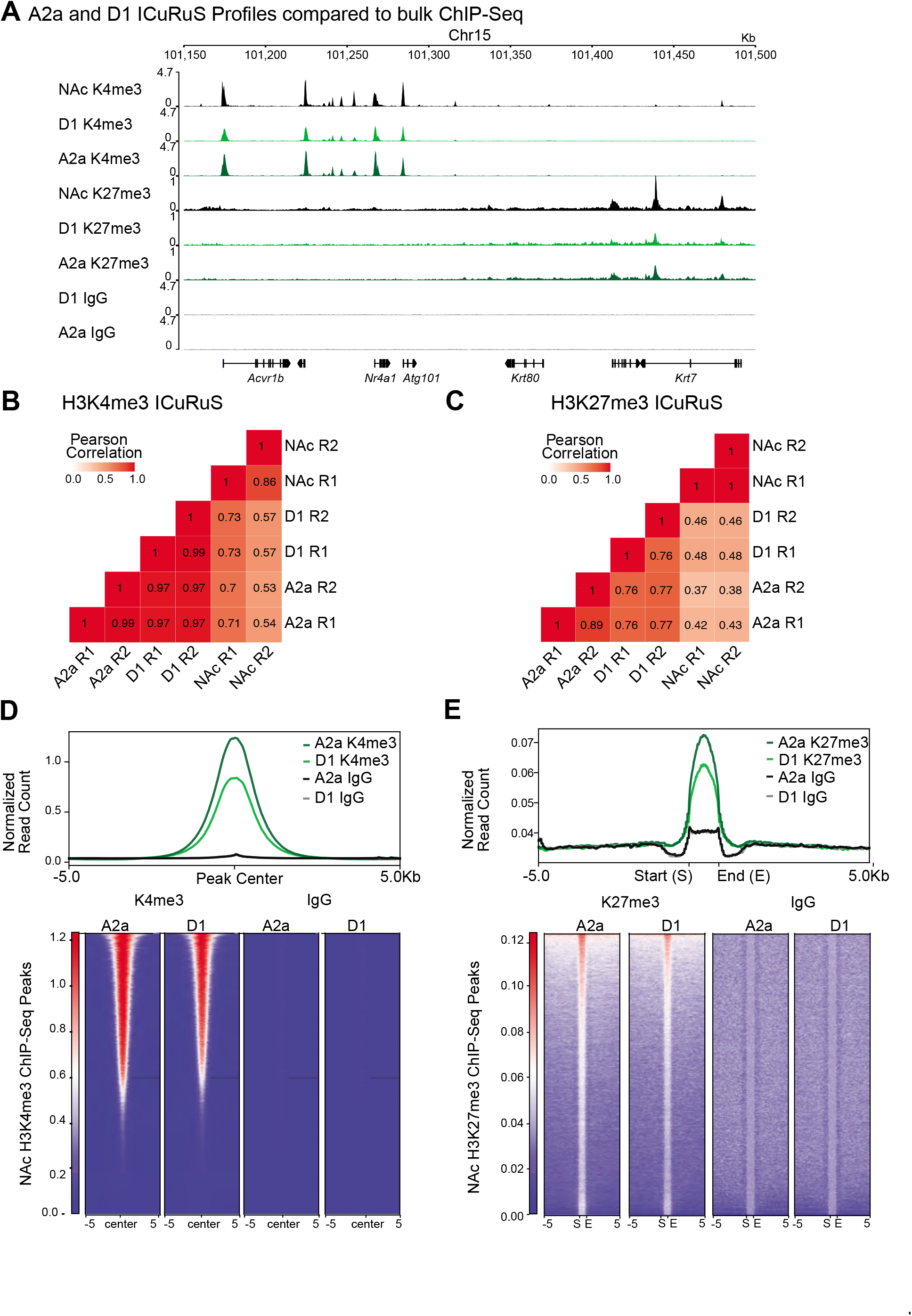
Epigenomic profiling in A2a and D1 neurons in the mouse striatum using ICuRuS. **A)** Representative genome browser views of different profiling methods and cell-types. Nucleus accumbens (NAc) bulk ChIP-seq data come from ^43^. Signal is normalized to total mapped reads: Count Per Million (CPM). **B)** H3K4me3 and **C)** H3K27me3 heatmap showing Pearson’s correlation coefficients among NAc ChIP-seq, D1 CnR, and A2a CnR. Two replicates are shown for each dataset. The correlation coefficients were calculated by dividing the genome into 1 kb bins and counting reads in each bin. **D)** Enrichment of A2a and D1 CnR H3K4me3 or IgG signal centered on peaks called from NAc H3K4me3 ChIP-seq by MACS2. Top 20,000 peaks sorted by MACS2 score were used. Heatmap shows signal around individual peaks, and the averaged signal is shown above the heatmap. Two replicates of CnR data were merged. Signal is normalized to total mapped reads: Count Per Million (CPM). **E)** Enrichment of A2a and D1 CnR H3K27me3 or IgG signal around peaks called from NAc H3K27me3 ChIP-seq by SICER. Peaks within 5kb were merged to avoid double counting. Heatmap shows signal around individual peaks, and the averaged signal is shown above the heatmap. Two replicates of CnR data were merged. Signal is normalized to total mapped reads: Count Per Million (CPM).

The accuracy and robustness of ICuRuS required considerable optimization. We found antibody selection to be the most important factor for successful ICuRuS (See Methods for antibody list). The same antibodies that produced robust data in ~10K N2a cells did not produce comparable data in A2a and D1 isolated nuclei (**Supplementary Figure 2A**). Specifically, use of H3K4me3 Antibody 3 (Ab3) or H3K27me3 Ab2 in N2a cells resulted in specific enrichment around peaks called from N2a H3K4me3 ChIP-seq data ^42^ **(Supplemental Figure 2C, E, F)** but nonspecific enrichment in A2a or D1 isolated nuclei **(Supplemental Figure 2K-P)**. These data highlight differences between in vitro and in vivo CnR that can be addressed by rigorous antibody selection when validating additional hPTMs beyond H3K4me3 and H3K27me3 in mouse brain.

### ICuRuS epigenomic profiling in A2a and D1 nuclei

To interrogate the relationships between cell-type specific hPTMs and mRNA expression, we integrated A2a and D1 H3K4me3 and H3K27me3 profiles with published mRNA profiles of D2- and D1-specific translatosomes (i.e. translating RNA profiling by Ribo-Tag) ^19^. As expected, A2a and D1 showed cell-type specific enrichment of H3K4me3 or H3K27me3 profiles at *A2a* and *Drd1*. Specifically, A2a nuclei were enriched with H3K4me3 and depleted of H3K27me3 at *A2a*, relative to D1 nuclei, which correlates with cell-type specific expression of A2a mRNA (**Figure 3A**). D1 nuclei were enriched with H3K4me3 and depleted of H3K27me3 at *Drd1*, relative to A2a expressing nuclei, which correlates with cell-type specific expression of D1 mRNA **(Figure 3B)**. In contrast, *CamK2a*, which is expressed in both cell types, was enriched with H3K4me3 and depleted of H3K27me3 in both A2a and D1 nuclei **(Figure 3C)**. Next, we quantified the association between overall patterns of cell-type specific gene expression and H3K4me3 or H3K27me3 enrichment. To accomplish this, we segmented H3K4me3 and H3K27me3 signal across three gene expression groups in A2a and D1 nuclei: high, (Fragments Per Kilobase Million, FPKM >= 10), medium (1 <= FPKM < 10) and low gene expression (FPKM < 1) ^19^. A2a H3K4me3 reads were enriched in the promoter of highly expressed A2a genes **(Figure 3D)**and A2a H3K27me3 reads were enriched in lowly expressed genes **(Figure 3E)**. Similarly, D1 H3K4me3 reads were enriched in the promoter of highly expressed D1 genes **(Figure 3F)**and D1 H3K27me3 reads were enriched in lowly expressed genes **(Figure 3G)**. These results were consistent with the known association of H3K4me3 with gene activation and H3K27me3 with gene repression ^32^ and showed that H3K4me3 and H3K27me3 ICuRuS profiles corresponded with predicted hPTM enrichment based on cell-type specific gene expression. Finally, we observed that both global H3K4me3 **(Supplemental Figures 3E, F;** t_(19999)_ = −131.01, p < 2.2e-16) and global H3K27me3 **(Supplemental Figure G, H;** t_(40405)_ = −86.189, p < 2.2e-16) signal in peaks were greater in A2a than D1, although, the majority of H3K4me3 and H3K27me3 peaks were observed in both A2a and D1 **(Supplemental Figure 3A, B)**.

**Figure 3.**
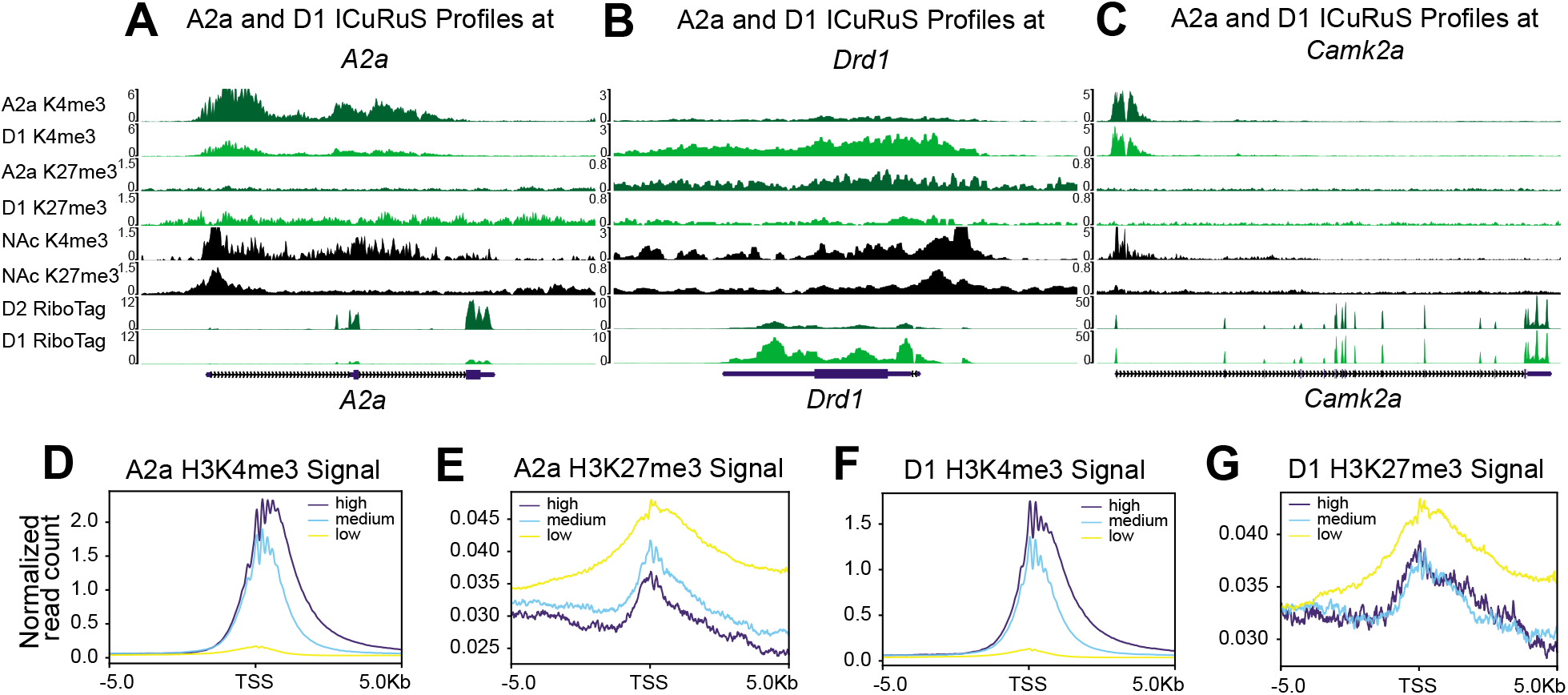
Comparison of peak loci across cell-types. **A-C)** A2a and D1 specific H3K4me3 and H3K27me3 modifications around known cell-type markers *A2a* **(A)**, *Drd1* **(B)**, and *Camk2a* **(C)**. **D**) A2a H3K4me3 signal, (**E)** A2a H3K27me3 signal, **(F)** D1 H3K4me3 signal, **(G)** and D1 H3K27me3 signal centered on transcription start site (TSS). For each cell-type and hPTM, genes are categorized into 3 groups: high (FPKM >= 10), medium (1 <= FPKM < 10), and low (FPKM < 1) based on published MSN-specific Ribo-Tag ^19^. Signal is normalized to total mapped reads: Count Per Million (CPM).

### ICuRuS identified H3K4me3 and H3K27me3 co-enriched promotors in A2a and D1 nuclei

To further our understanding of the combinatorial function of hPTMs in regulating gene expression, we next examined A2A and D1 gene expression as a function of K27me3/K4me3 co-enrichment ^32^ **(Figure 4A, B)**. Beyond the binary states of active or repressed gene expression, genes co-enriched with H3K4me3 and H3K27me3 have been described as ‘poised,’ such that expression is low at baseline but high upon stimulation ^30^. To investigate this concept in specific MSNs, we defined bivalency as coincident H3K4me3 and H3K27me3 enrichment between 2 kb upstream and 1 kb downstream of the TSS, in A2a and D1 nuclei ^44^. ICuRuS found that co-enriched H3K4me3 and H3K27me3 genes are expressed at an intermediate level relative to H3K4me3- or H3K27me3-alone (Wilcoxon rank sum test with Benjamini-Hochberg correction, P < 2e-16**; Figure 4A, B)**. Next, we calculated the global mutually exclusive index as the random vs real coincidence of two measured hPTMs ^45^. We found the K4me3/K27me3 mutually exclusive index is greater in D1 than A2a nuclei (**Figure 4C**), indicating that one or both hPTMs occurs less often in the TSS of D1 MSNs when compared to A2a MSNs (**Figure 4C**). However, among the four hPTM profiles analyzed, A2a, D1 and N2a were above 1 and greater than NAc which suggests A2a and D1 ICuRuS reveals distinct features not observed in NAc bulk analysis (**Figure 2C**). As expected, the majority of co-enriched domains are common to both A2a and D1 MSNs (**Figure 4D**) and enriched for GO biological processes such as negative regulation of cell differentiation, nervous system development and extracellular matrix organization (**Figure 4E**).

**Figure 4.**
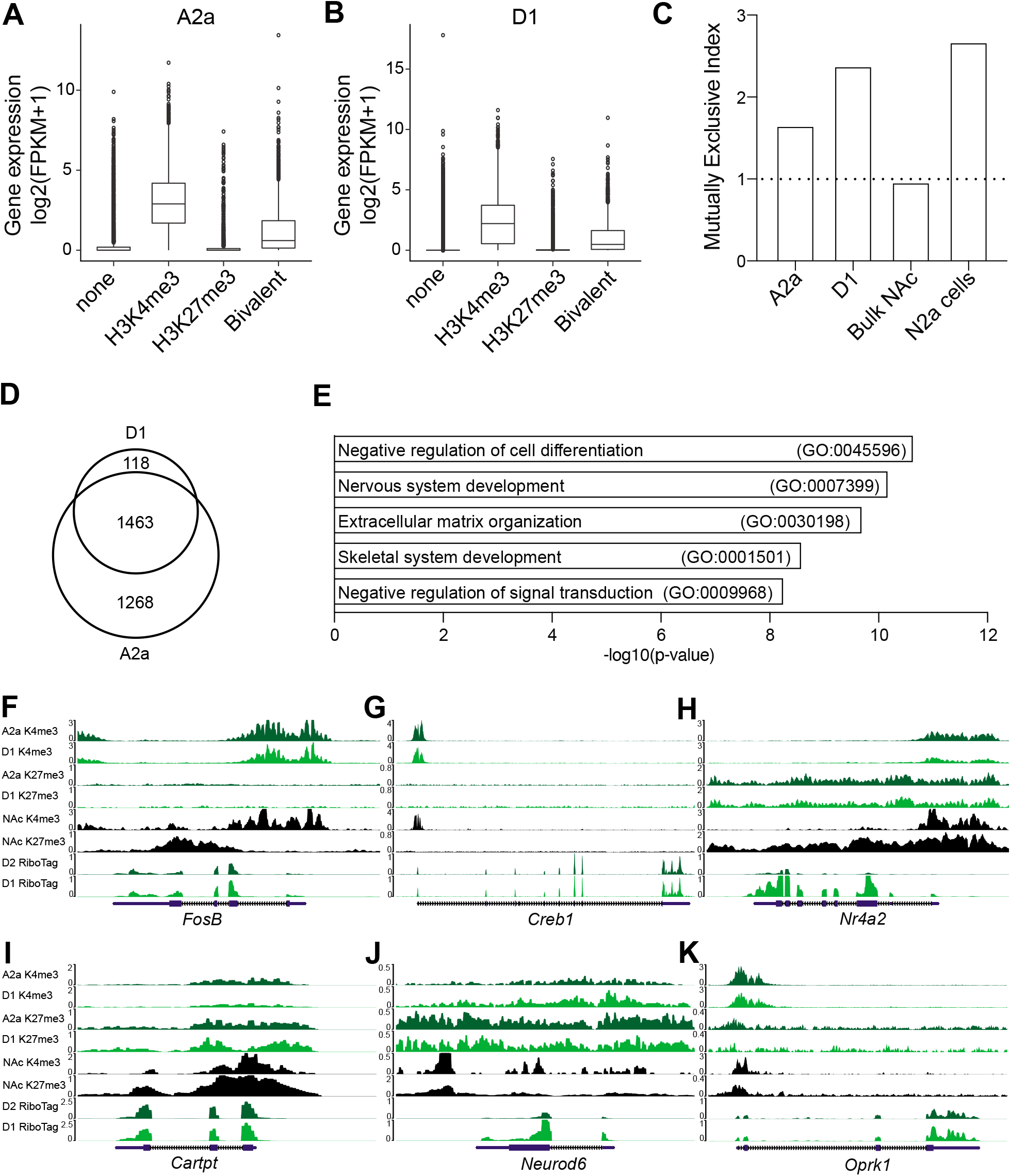
ICuRuS identified H3K4me3 and H3K27me3 co-enriched promotors in A2a and D1 nuclei. **A)** A2a and **(B)** D1 H3K4me3, H3K27me3, and H3K4me3/H3K27me3 combinatorial enrichment between 2 kb upstream and 1 kb downstream of the promoter start site TSS for cell-type specific genes using published MSN-specific Ribo-Tag^19^. **C)** H3K4me3/H3K27me3 mutually exclusive index for A2a, D1, N2a, and bulk NAc (bulk NAc raw data was extracted from ^43^). A dashed line with an index equal to 1.0 indicates that no exclusive effects exist. **D)** Venn diagram displaying cell-type specific and overlapping bivalent domains in A2a and D1 nuclei. **E)** GO biological processes for the top five bivalent domains that are present in both A2a and D1 cell-types. **F-K)** Genome browser view of **(F)** *FosB,* **(G)** *Creb1*, **(H)** *Nr4a2*, **(I)** *Cartpt*, **(J)** *Neurod6*, and **(K)** *Oprk1* for different profiling methods and cell-types. Nucleus accumbens (NAc) bulk ChIP-seq data come from ^43^. Ribotag data come from^19^. Signal is normalized to total mapped reads: Count Per Million (CPM).

Finally, we profiled K4me3/K27me3 co-enrichment at genes at immediate early genes (IEGs) that show rapid activation in response to stimulation and are expressed in both cell-types. IEGs *Nr4a1*, *Fosb*, and *Creb1*, are enriched in H3K4me3 but not H3K27me3 in both A2a and D1 nuclei (**Figure 2A, Figure 4F, G, respectively**). In contrast, *Nr4a2,* which is expressed more in D1 than A2a neurons, and *Cartpt,* which is A2a specific ^46,47^, were both K4me3/K27me3 co-enriched in both cell-types (**Figure 4H, I**, **Supplementary Figure 4A**). The profiles of *Nr4a2* and *Cartpt* suggested that additional hPTMs may confer D1 or A2a specificity, respectively (**Figure 4I**). We found D1-specific gene, *Neurod6*, was K4me3/K27me3 co-enriched in D1 nuclei and enriched only in H3K27me3 in A2a nuclei (**Figure 4J**), while D1-specific gene, *Oprk1*, was co-enriched in A2a nuclei and depleted in H3K27me3 in D1 nuclei (**Figure 4K**). In contrast, *Bdnf* is expressed higher in D1 than A2a nuclei but showed similar co-enrichment of H3K4me3 and H3K27me3 in both A2a and D1 nuclei (data not shown). Taken together, combinatorial hPTM profiling by ICuRuS suggested that A2a and D1 co-enriched genes are likely suppressed in a cell-type specific manner during development but retain some level of gene activity ^33,48,49^.

## Discussion

In developing the hybrid ICuRuS protocol, we sought to answer the following question: how do subtle changes in chromatin orchestrate cell-type specific functions that produce holistic changes in phenotype? We found that ICuRuS generated robust and reproducible H3K4me3 and H3K27me3 profiles at sufficient depth to examine cell-type specific chromatin from a single mouse striatum. These findings are an improvement over current methods, such as bulk tissue CnR^7,40^, which obscures cell-type specificity, and single-cell CnR, which requires pooled brain tissue and obscures subject variability ^4,5^, a critical barrier given individual differences in stimulus-induced gene expression and susceptibility to psychiatric disease ^33,50–52^. Recent studies have applied FACS to isolate MSNs, but we found that the affinity purification strategy used in ICuRuS (and in the original INTACT protocol ^9^) potentially avoids FACS induced ectopic upregulation of activity dependent genes ^53–55^ . Finally, ICuRuS allowed identification of K4me3/K27me3 co-enriched domains at cell-type specific genes, supporting the hypothesis that neuronal subtype specific gene expression is regulated via combinatorial hPTMs.

In the mouse striatum, A2a and D1-MSNs have complimentary, opposing, or restricted roles in neuronal function ^59–64^. A2a and D1 MSNs make up ~95% of the neuronal population in the striatum ^65,66^. Cell type specific gene expression studies have genetically define these subpopulations ^19,46,47,67^, reporting that distinct transcriptional networks in D1-MSNs promote the motivation to seek drugs, while A2a-MSNs generally inhibit this behavior in mice ^68,69^. Conversely, depending on the temporality of stimulation, activation of A2a-MSNs can enhance motivation ^70–72^. Our previous work shows that *Nr4a1* activation in either A2a alone (unpublished) or in bulk striatal neurons in mouse, reduces the motivation to seek cocaine following abstinence ^36^. In whole ventral striatum, Nr4a1 binds and activates downstream target gene, *Cartpt*, via depletion of H3K27me3 and enrichment of H3K4me3 at this locus ^36^. We find here using ICuRuS that H3K27me3 is lowly enriched at *Nr4a1* in A2a and D1 nuclei, consistent with strong activation upon cocaine exposure ^36^, while *Cartpt* is enriched in both H3K4me3 and H3K27me3 in both A2a and D1 nuclei. Interestingly, H3K4me3 was also enriched at several *Nr4a1* exons suggesting alternative transcription start sites ^73^. We posit that activation of *Nr4a1* at alternative promoters may produce Nr4a1 isoforms that lack the N-terminal transactivation domain, which has important implications on downstream target gene activation ^74^. In addition, cell-type specific H3K27me3 depletion at *Cartpt* could establish the mechanism that confers A2a specific expression of this peptide. Our lab is adequately equipped to address the causal relevance of these findings using cell-type specific CRISPR activation (e.g. dCas9-VPR) targeted to alternative *Nr4a1* promoters and CRISPR epigenetic editing of H3K27me3 using K27me3-specific demethylase, Friend of GATA1 (FOG1) at *Cartpt* ^75^. Future studies will use cell-type specific epigenetic editing to determine the exact function of hPTMs at gene loci and how cell-type specific hPTM enrichment changes in response to stimuli.

Traditionally, H3K4me3 and H3K27me3 co-enriched genes have been described as ‘poised’ – simultaneously silenced but responsive to stimuli ^32,76,77^. Consistent with the literature, MSN ICuRuS profiling showed binary regulation of gene expression (on/off) when H3K4me3 and H3K27me3 were independently enriched. However, genes co-enriched in H3K4me3 and H3K27me3 showed intermediate levels of gene expression across both MSN subtypes. These data suggest competitive antagonism between the transcription factors recruited by H3K4me3 and H3K27me3. In this model, the expression level of specific transcription factors may explain discordance in binary gene regulation ^78,79^. Intriguingly, H3K4me3 and H3K27me3 co-enrichment largely defines developmental genes ^80^, and priming events that regulate delayed changes in gene expression ^36,77,81–83^. While ICuRuS can determine K4me3/K27me3 co-enrichment at specific loci, it cannot determine if these marks occur on the same histone or nucleosome. Currently our lab is investigating if K4me3/K27me3 co-enriched loci show true bivalency using sequential ChIP in mouse striatum ^48,84–87^.

While consideration of combinatorial hPTM function is crucial to interpreting the histone code, current data is sparse on the function of bivalent chromatin in the adult mammalian brain or in response to environmental stimuli ^81^. IEGs *Fosb, Creb1,* and *Nr4a1* lack broad H3K27me3 domains in A2a and D1 nuclei ^88–91^, suggesting that additional repressive hPTMs are likely involved in the poised state of *Fosb, Creb1,* and *Nr4a1*. Whereas *Fosb, Creb1,* and *Nr4a1* show no MSN-subtype specific expression or hPTM enrichment*, Cartpt, Neurod6, Nr4a2, Oprk1 and Bdnf* are differentially expressed between D1 and A2a cells ^19,46,47,67,92^ and showed differential H3K7me3 and H3K4me3 enrichment in either A2a or D1. For example, D1 specific expression of *Neurod6* was associated with the relative depletion of H3K4me3 D1 MSNs ^19,46,92^. A2a specific expression of *Oprk1* was associated with depletion of H3K27me3 in A2a cells ^19,46,92^. In this way, neuronal subtype specific expression is conferred by either loss of activating hPTMs in the non-expressing subtype, or loss of a repressive hPTM in the expressing subtype. Alternatively, A2a-specific *Cartpt* expression and D1-specific *Bdnf* and *Nr4a2* expression were not associated with cell-type specific differences in H3K4me3 or H3K27me3 enrichment, suggesting that additional components of the histone code are at play. Taken together, we conclude that K4me3/K27me3 co-enrichment was associated with a subset of genes expressed in a cell-type specific manner. Elucidating the epigenetic mechanisms that govern MSN subtype-specific gene expression has emphasized the role of combinatorial hPTMs, as a key determinant of cell-type specificity. Taken together, cell-type specific hPTM profiling may reveal the mechanisms by which environmental stimuli activate gene expression in only one neuronal subtype. Defining cell-type specific hPTM profiles will have important implications on neuropathology ^91^.

ICuRuS serves as highly adaptable method for investigations into cell-type specific epigenetic gene regulation in brain. First, it provides high resolution hPTM profiles from low cell numbers achievable from a single mouse at relatively low cost. Published expression profiling data defining specific cell-types of investigators can easily target any cell-type in the brain using CRE transgenic mice or viruses for the expression of SUN1-GFP for target cell immunoprecipitation^8,67^. Second, we identified hPTM enrichment patterns that can be manipulated using cell-type specific epigenetic editing tools to establish causal roles of hPTMs in the regulation of gene expression and behavior ^40,93–95^. Finally, ICuRuS can be used to casually link specific hPTM to gene activity in response to environmental stimuli. Altogether, our findings complement and expand upon existing methods for the examination of cell-type specific global chromatin changes in brain.

## Methods

### Animals

The *R26-CAG-LSL-Sun1-sfGFP* knock-in mouse on the C57BL/6J background was crossed with A2a-cre and Drd1-cre mice C57BL/6J background to generate Sun1-sfGFP;A2a-cre and Sun1-sfGFP;Drd1-cre mouse lines. Mice were housed on a 12-h light-dark cycle at constant temperature (23 °C) with access to food and water ad libitum. All animal procedures were conducted in accordance with the National Institutes of Health Guidelines as well as the Association for Assessment and Accreditation of Laboratory Animal Care. Ethical and experimental considerations were approved by the Institutional Animal Care and Use Committee of The University of Pennsylvania.

### INTACT

For each experiment, bilateral striatum of mouse was rapidly dissected in ice-cold homogenization buffer (0.25M sucrose, 25mM KCl, 5mM MgCl2, 20mM Tricine-NaOH). The tissue was the slow frozen at −80C until INTACT was conducted. The INTACT procedure used was modified rom Mo et al., 2015. To initiate the INTACT procedure, tissue was Dounce homogenized using a loose pestle (10 strokes) in 1.2 mL of homogenization buffer supplemented with 1mM DTT, 0.15mM spermine, 0.5mM spermidine, and EDTA-free protease inhibitor (Roche). A 5% IGEPAL-630 solution was added, and the homogenate was further homogenized with the tight pestle (7 strokes). The sample was then mixed with 1.3 mL of 50% iodixanol density medium (Sigma D1556), and added to an Ultra-Clear Tube (13.2mL, Beckman and Coulter, 344059). The sample was then underlayed with a gradient of 30& and 40& iodixanol, and ultra-centrifuged at 13,000rpm for 18 minutes in a swinging bucket centrifuge at 4°C. Nuclei were collected at the 30&-40& interface and pre-cleared by incubating with 20 μL of Protein G Dynabeads (Life Technologies 10003D) for 15 minutes. After removing the beads with a magnet, the mixture was incubated with 10 μL of 0.2 mg/mL rabbit monoclonal anti-GFP antibody (Life Technologies G10362) for 30 minutes. 60 μL of Dynabeads were added, and the mixture was incubated for an additional 20 minutes. To increase yield, the bead-nuclei mixture was placed on a magnet for 30 seconds to 1 minute, completely resuspended by inversion, and placed back on the magnet. This was repeated seven times. Bead-bound nuclei were then washed 3 × 800 uL with wash buffer. All steps were performed on ice or in the cold room, and all incubations were carried out using an end-to-end rotator. When purifying RNA following INTACT, RNasin ® Plus Ribonuclease Inhibitor (Promega N2611) was added to the antibody and bead buffers.

### Cell Counting

For cell counting to quantify INTACT specificity and yield, during the INTACT procedure aliquots were collected following nuclei isolation and following affinity bead-bound pull downs. Nuclei from isolation (i.e. before the addition of antibody and Protein G beads) and bead-bound nuclei were stained with DAPI (30uM) and 10ul of sample was added to a hemocytometer. Images were acquired using a Leica Fluorescence Microscope. DAPI+/GFP+ and DAPI+/GFP-nuclei were counted using a hemocytometer. The following equations (Mo et al., 2015) were used to calculate the specificity (A) and yield (B) from INTACT:

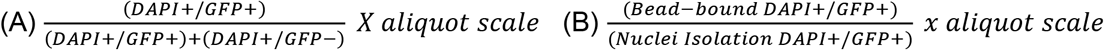

### CnR (following INTACT)

As the INTACT protocol relies on bead-immunopurification of Sun1-GFP+ nuclei, it is ideally suited to downstream processing of bead-immobilized nuclei by CnR. CnR was carried out according to published protocols (Skene & Henikoff, 2017), with changes pertinent to neuronal isolation. Nuclei from one striatum (~10,000) were washed 2X and then resuspended in Digitonin Buffer with either anti-H3K4me3 (<u>Antibody 1</u>: Abcam, Ab8580 (Figure 2); <u>Antibody 2</u>: Active Motif, 39159 (Supplemental Figure 2); <u>Antibody 3</u>: Epicyphr, 13-0041), anti-H3K27me3 (<u>Antibody 1</u>: Active Motif, 39055; <u>Antibody 2</u>: Thermo Fisher, MA5-1198) or IgG (Epicypher, 13-0042) for two hours, 4°C with rotation. Following incubation, bead-bound nuclei were washed 2X with Digitonin Buffer, and antibody-enriched fragments were cut by one-hour, 4°C incubation with 0.5 ul MNAse (20x CUTANA™ pAG-MNase 15-1116). Following MNase incubation, bead-bound nuclei were washed 2X with Digitonin Buffer, and antibody-enriched fragments were released by two-hour, 4°C incubation with 100mM CaCl_2_. Antibody-enriched fragments from bead-bound nuclei were then washed again and the reaction was stopped with 33 ul Stop Buffer (68 ul of 5M NaCl, 40 ul of 0.5M EDTA, 40 ul of 4mM EGTA) and incubated for 10 min (37°C). 1 μl of 10% (wt/vol) SDS and 1.5 μl of proteinase K (20 mg/ml) were added to each sample, followed by 20 min (50 °C) incubation. Total DNA from each sample was purified using phenol/chloroform/isoamyl alcohol and dissolved in TE.

### Library Prep

Library preparation of CnR was performed using a NEB Ultra II library preparation kit, with modifications to the protocol ^39^. CnR DNA (<5ng) was used as input. End preparation was performed for 30 minutes at 20°C followed by 1 hour at 50°C. Adapter at .6 pmol was ligated to end preparation products at 20°C for 15 minutes. USER enzyme was added, and DNA was purified using 1.1X AMPure beads. Libraries were amplified with 2x Ultra Q5 min, universal primer, and index primers. PCR was as follows: 98°C for 20s; 2 cycles of 98 °C for 10 s and 65 °C for 10 s; and a final extension at 65 °C for 5 min. PCR products were size selected using 1.1X AMPure beads. PCR products were quantified using quBIT, Agilent Bioanalyzer and NEB library quant kit (E7630L). Libraries were pooled at equimolar amounts and sequenced using the Nextseq500 platform. Paired-end sequencing was performed (read length, 38 bp × 2; index, 8 bp).

### Immunohistochemistry

Sun1-GFP; A2a-cre and Sun1-GFP; Drd1-cre mice were anesthetized with ketamine/xylazine and perfused with 4% paraformaldehyde (PFA). Brains were then rapidly extracted and stored in 4% PFA overnight for fixation. Brains were then transferred to 15% sucrose for 24 hours and then stored in 30% sucrose until sectioning (40 μm) using a cryostat. Sections were blocked for 90 minutes at room temperature with 10% Normal Donkey Serum and 0.2% Triton X-100 in PBS and then incubated with 0.2% Triton X-100 in PBS with the following antibodies overnight at 4°C: rabbit anti-Drd1 (1:500, Bioss USA BSM-52920R) or rabbit anti-A2a (1:200, Fisher Scientific PA1-042), co-incubated with goat anti-GFP (1:500, Rockland, 600-101-215). Sections were then washed three times with PBS and incubated at room temperature in the dark for two hours with secondary antibodies for fluorescent labeling: Donkey anti-Rabbit IgG (H+L) Highly Cross-Adsorbed Secondary Antibody, Alexa Fluor Plus 555 (Fisher Scientific A32794) and Donkey anti-Goat IgG (H+L) Cross-Adsorbed Secondary Antibody, Alexa Fluor 488 (Fisher Scientific A-11055). After the first hour of incubation, 0.5 μg/mL of DAPI (dihydrochloride, Invitrogen D1306) was directly added into the secondary antibody solution. Sections were then washed three times with PBS and mounted on glass slides using ProLong™ Gold Antifade Mountant (Invitrogen P36930). Images were taken using a Zeiss LSM 810 Confocal Microscope.

### Published ChIP-seq data analysis

N2a and nucleus accumbens (NAc) ChIP-seq data were previously published (H3K4me3 ^42^; H3K27me3 ^41^; NAc H3K4me3 and H3K27me3^43^). Raw reads were aligned to the mm10 reference genome using bowtie2 (Langmead and Salzberg, 2012, PMID: 22388286) with parameters: -q -- no-unal --phred33. Uniquely mapped reads (mapping quality score >= 20) were selected using samtools with command samtools view -q 20 (version 0.1.19) ^96^.

### CnR data processing and analysis

Base call (BCL) files were demultiplexed and converted into FASTQ files using bcl2fastq2 (version v2.20.0.422) with default parameters. Next, raw paired-end reads were mapped to the mm10 reference genome using bowtie2 (version 2.1.0)^97^ with options: -q --local --no-mixed --no-unal -- dovetail --phred33. Using local alignment mode makes removal of adapter sequences unnecessary. Uniquely mapped reads (mapping quality score >= 20) were selected using samtools (version 0.1.19, samtools view -q 20). Then, we used Picard (version: 2.23.4)^98^ to check whether the insert size distribution is consistent with the library fragment size obtained from Bioanaylzer. Next, duplicates were removed using Picard MarksDuplicates function. We selected deduplicated mapped reads with insert size between 150 and 500 bp using samtools for downstream analyses. Read alignments were normalized to total mapped reads using deepTools with command: bamCoverage –bam inbam -o outbw –normalizaUsing CPM and visualized by deepTools pyGenomeTracks (https://github.com/deeptools/pyGenomeTracks).

### RNA-seq data processing and analysis

Drd1 and A2a cell-type-specific RNA-seq data were derived from RiboTag-purified Drd1 and Drd2 neurons of the NAc, respectively ^19^. Raw reads were mapped to the mm10 reference genome and Gencode annotation (vM23) using STAR ^99^with parameters: --outFilterMismatchNmax 3 -- outFilterMultimapNmax 1 --alignSJoverhangMin 8. Differentially expressed genes between Drd1 and Drd2 neurons were detected using cufflinks cuffdiff with default parameters (v2.2.0) ^100^. Read alignments were normalized to total mapped reads using deepTools for visualization.

### Correlation between ICuRuS and ChIP-seq data

To evaluate reproducibility and specificity of the ICuRuS data, we calculated pairwise Pearson’s correlation coefficients among ICuRuS replicates and corresponding ChIP-seq datasets. We first computed with read coverages for the entire genome using deepTools with multiBamSummary function. Then, correlation coefficients were calculated and visualized by plotCorrelation function of deepTools ^101^ with parameters: --skipZeros.

### Signal around peaks or TSS

For signal around H3K4me3 peaks and TSS (+/− 5 kb), we calculated signal using deepTools with command: computeMatrix reference-point -a 5000 -b 5000. For signal around H3K27me3 peaks, peaks 5kb apart were first merged using the bedtools merge ^102^ with parameter: -d 5000. Then, H3K27me3 signal were calculated using deepTools with command: computeMatrix scale-regions -a 5000 -b 5000. Signal around each peak or TSS and was plotted using deepTools plotHeatmap function. Averaged signal was plotted using deepTools plotProfile function.

### Peak calling

H3K4me3 peaks were called using MACS2 ^103^ with parameters: -f BAM -g mm -B --keep-dup all -q 0.01. H3K27me3 peaks were called using SICER ^104^ with the command: sicer -t inbam -c control -s mm10. H3K27me3 peaks with false discovery rate less than 0.01 were selected. Overlapped peaks were counted using HOMER mergePeaks function ^105^ and visualized by Upset plots ^106^. Fraction of reads in peaks was calculated using a custom Python script.

### Bivalency Analysis

We categorized genes into four groups based on whether there are H3K4me3 and/or H3K27me3 peaks on promotor regions (−2 kb to +1 kb): ^44^ 1) Bivalent genes which have both H3K4me3 and H3K27me3 peaks, 2) H3K4me3 genes which have only H3K4me3 peaks, 3) H3K27me3 genes which have only H3K27me3 peaks, and 4) none genes which have no peaks. Mutually exclusive index was calculated based on a previous method ^107^.

## Data availability

All raw and processed sequencing data generated in this study have been deposited to NCBI Gene Expression Omnibus under accession number: GSE193673 (https://www.ncbi.nlm.nih.gov/geo). All scripts are available at github (https://github.com/HellerLAbeats/ICuRuS).

## Acknowledgments

Financial support was provided by NIH-NIDA Avenir Director's Pioneer Award (E.A.H, DP1 DA044250), NIDA Research Project Grant (E.A.H, 1R01DA052465-01A1), Pilot Award from the Epigenetics Institute at the University of Pennsylvania, NIH Drug Abuse Dissertation Research Award (M.D.C. R36 DA050877), T32 Predoctoral Training Grant in Addiction (D.K.F. NIDA T32DA028874), and SynGAP Research Fund Postdoctoral Fellowship (S.Z.). We thank Dr. Yemin Lan for independently validating some findings present in this study. Figure 1C and Supplemental Figure 2A were created with Biorender.com.

---

**Supplementary Figure 1.**
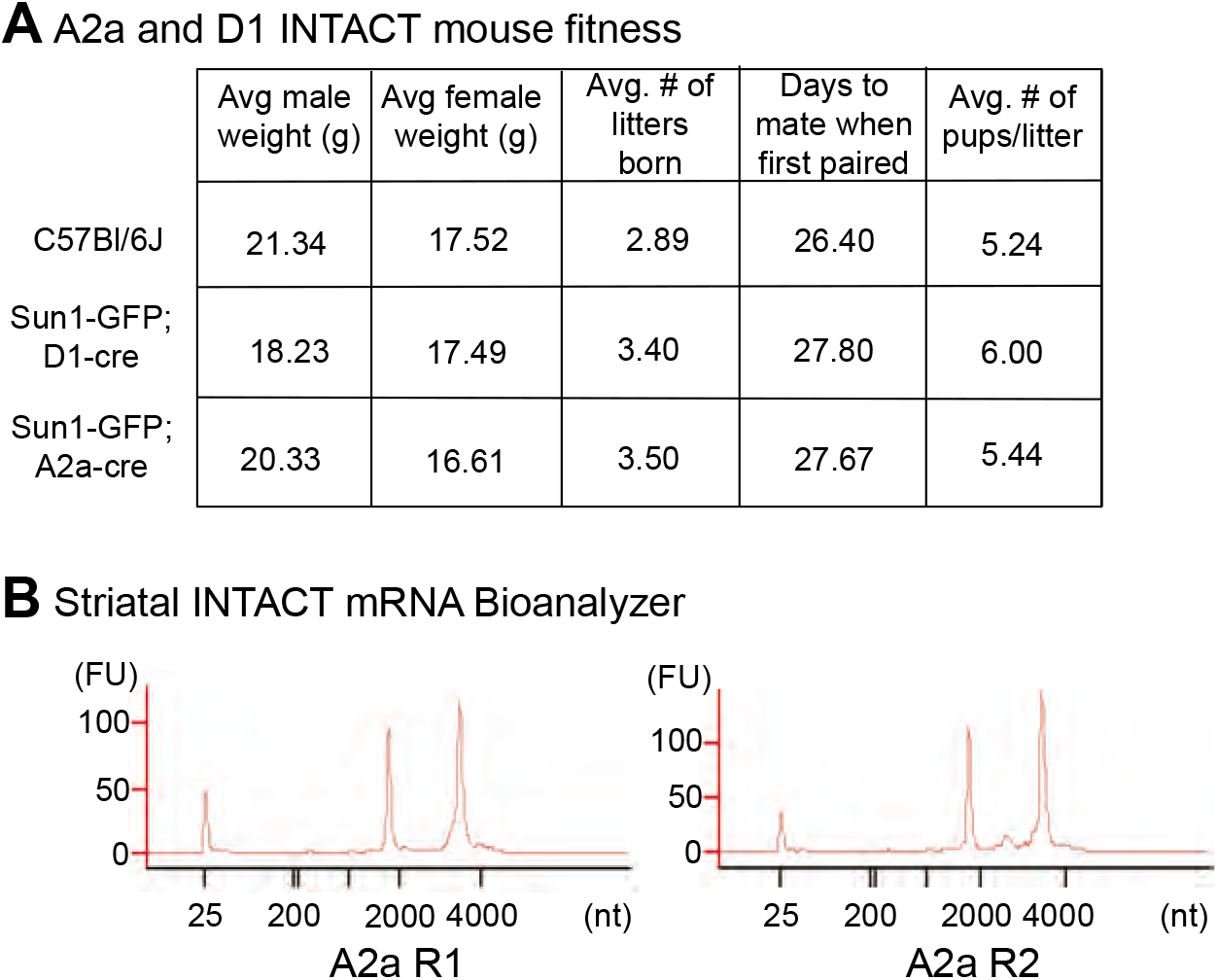
**A)** Animal weight and fitness data from Sun1-GFP; D1-cre and Sun1-GFP; A2a-cre mouse lines (P56-57 n = 6-8/group) **B)** Representative mRNA BioA outputs following A2a-INTACT using Agilent Pico RNA kit. RIN 9.80 and 9.70, respectively

**Supplementary Figure 2.**
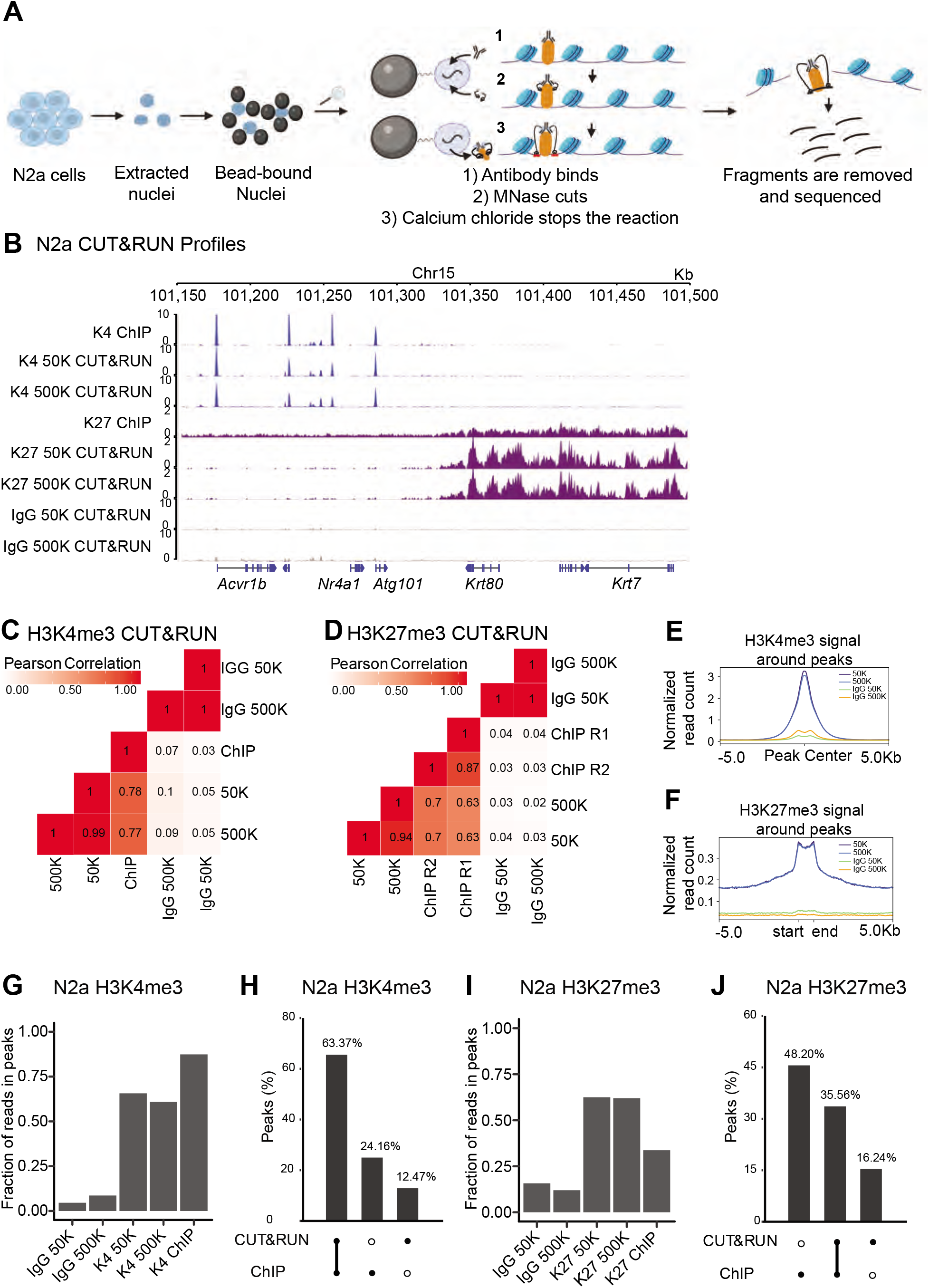
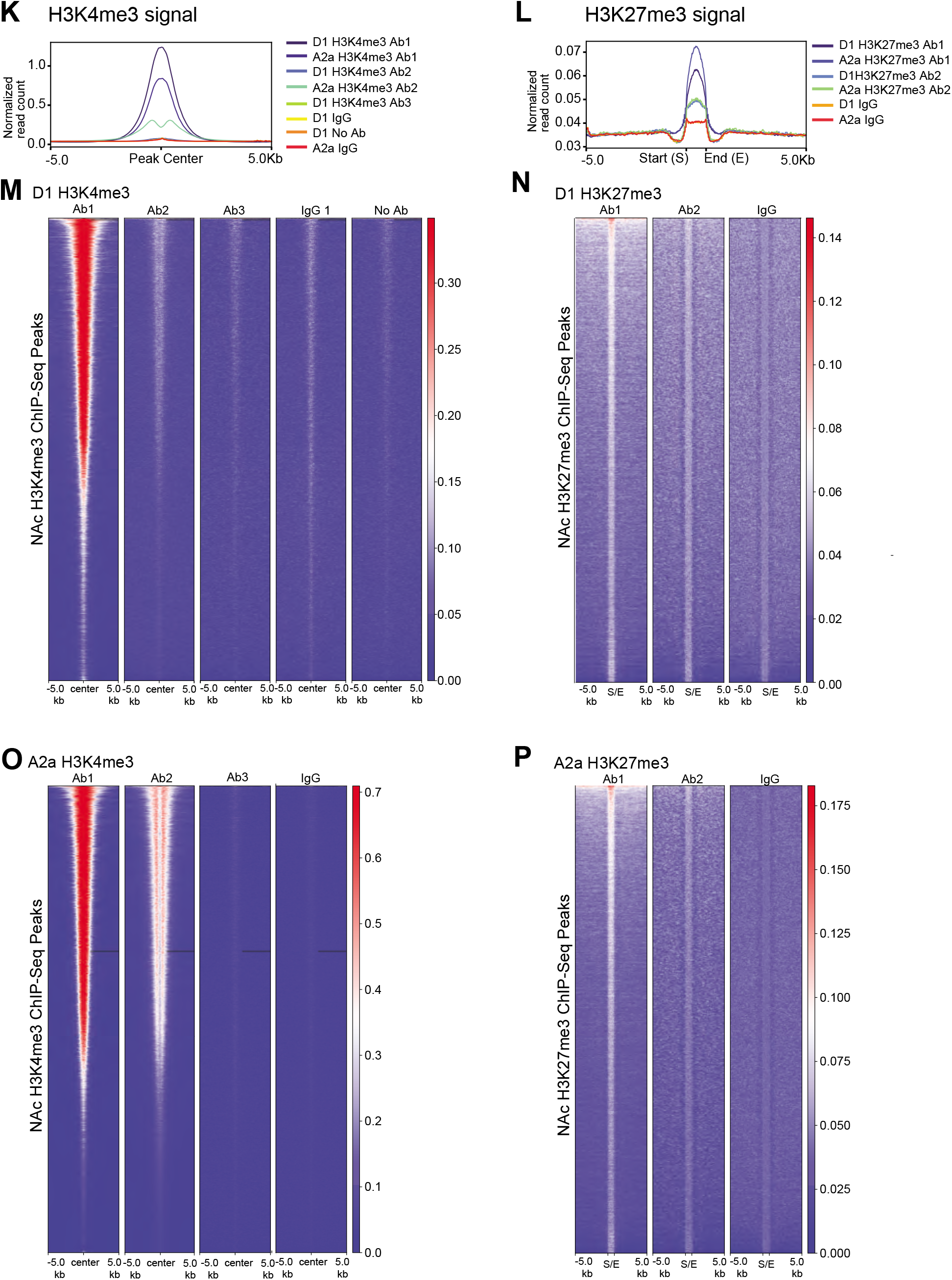
**A)** Schematic of CnR protocol on transfected N2a cells created at Biorender.com. **B)** Representative genome browser views of different profiling methods in N2a cells. Two N2a cell amounts 50,000 (50K) and 500,000 (500K) were tested for the CnR approach. Signal is normalized to total mapped reads: Count Per Million (CPM). N2a ChIP-seq data come from ^42^ (H3K4me3) and ^41^ (H3K27me3). **C)** H3K4me3 and **D)** H3K27me3 heatmap showing Pearson’s correlation coefficients among N2a ChIP-seq and CnR H3K4me3 signals, and N2a CnR IgG control. The correlation coefficients were calculated by dividing the genome into 2 kb bins and counting reads in each bin. **E)** Averaged N2a H3K4me3 or IgG signal centered on peaks called from N2a H3K4me3 ChIP-seq data by MACS2. **F)** Averaged N2a H3K27me3 or IgG signal around peaks called from N2a H3K27me3 ChIP-seq data by SICER. Peaks within 5kb were merged in order to avoid double counting. **G)** Fraction of reads in peaks called from N2a H3K4me3 ChIP-seq data for each indicated dataset. **H)** Overlapped and individual peaks between N2a ChIP-seq and CnR H3K4me3 data sets. **I)** Fraction of reads in peaks called from N2a H3K27me3 ChIP-seq data for each indicated dataset. **J)** Overlapped and individual peaks between N2a ChIP-seq and CnR H3K27me3 data sets. **K-L)** Density plots of different **(K)** H3K4me3 and **(L)** H3K27me3 antibodies used for A2a- and D1-specific ICuRuS. H3K4me3 Antibody 1 (Ab1): Abcam; H3K4me3 Antibody 2 (Ab2): Active Motif; H3K4me3 Antibody 3 (Ab3): Epicyphr; H3K27me3 Antibody 1 (Ab1): Active Motif; H3K27me3 Antibody 2 (Ab2): Thermo Fisher. **M-N)** Heatmaps from different **(M)** H3K4me3 and **(N)** H3K27me3 antibodies D1-specific ICuRuS. **O-P)** Heatmaps from different **(O)** H3K4me3 and **(P)** H3K27me3 antibodies A2a-specific ICuRuS.

**Supplementary Figure 3.**
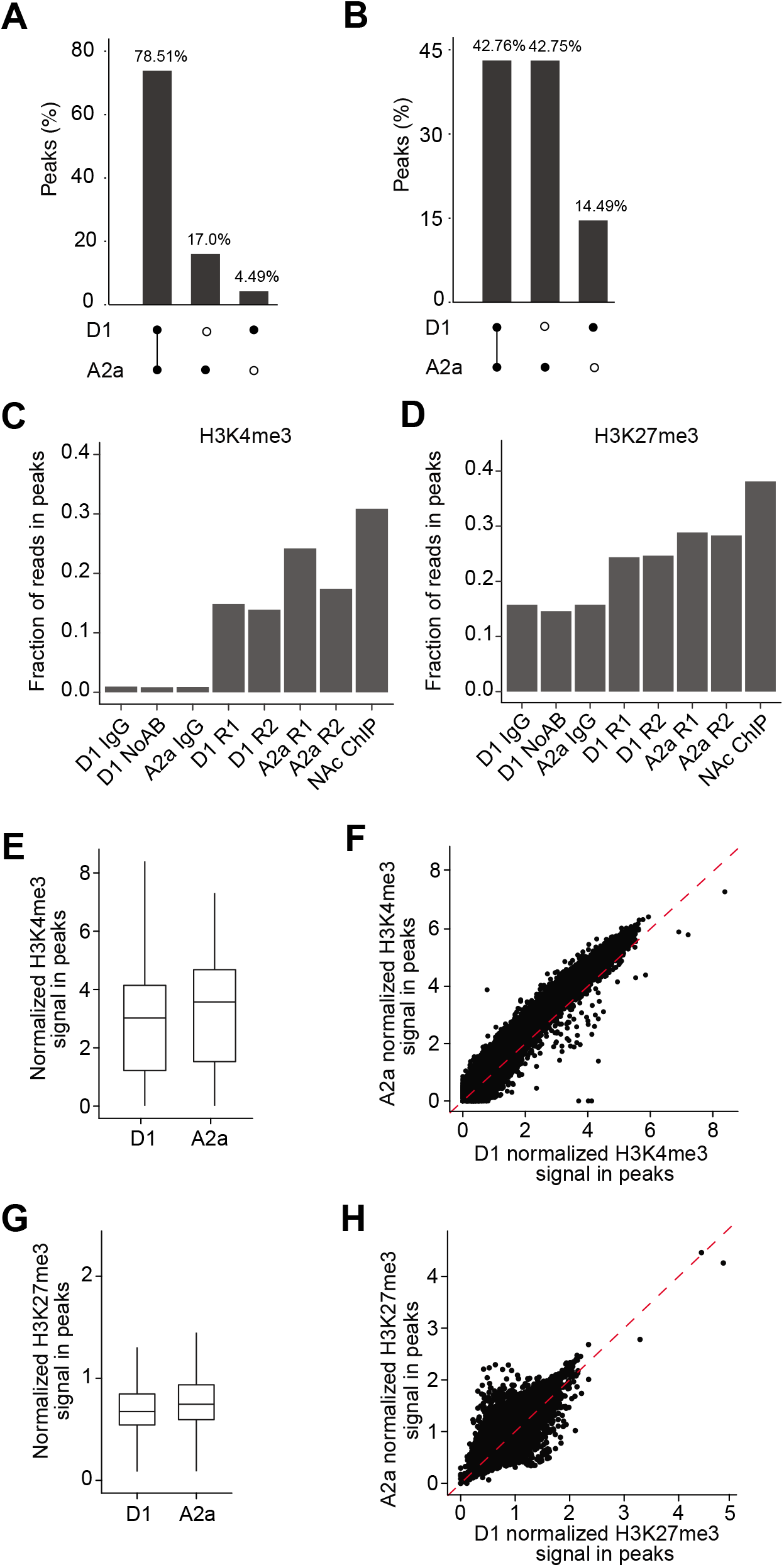
**A-B)** Overlapped and individualized peaks between A2a and D1 **(A)** H3K4me3 and **(B)** H3K27me3 ICuRuS. **C-D)** Fraction of reads in peaks called from NAc **(C)** H3K4me3 and **(D)** H3K27me3 ChIP-seq data for each indicated dataset. Nucleus accumbens (NAc) bulk ChIP-seq data come from ^43^. Boxplot **(E)** and scatterplot **(F)** showing H3K4me3 signal in peaks called from NAc H3K4me3 ChIP-seq data in A2a and D1 ICuRuS. Signal is normalized to total mapped reads: Count Per Million (CPM). Boxplot **(G)** and scatterplot **(H)** showing H3K27me3 signal in peaks called from NAc H3K27me3 ChIP-seq data in A2a and D1 ICuRuS. Signal is normalized to total mapped reads: Count Per Million (CPM).

**Supplementary Figure 4.**
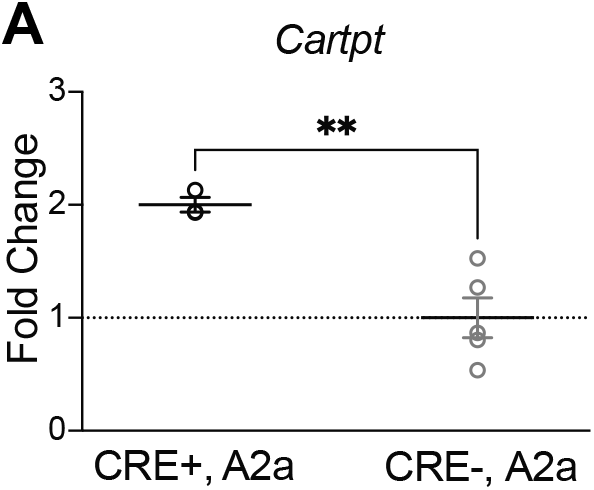
**A)** qPCR following A2a-INTACT for A2a-INTACT Cre+ vs A2a-INTACT Cre- (bulk supernatant) for *Cartpt* revealed *Cartpt* expression is increased in A2a+ nuclei compared to bulk supernatant (t_(6)_ = 4.183, p = 0.0058).

**Supplementary File 1.** A2a and D1 ICuRus Peaks

## Notes

### Competing Interest Statement

The authors have declared no competing interest.

